# Early divergence of mutational mechanisms drives genetic heterogeneity of fetal tissues

**DOI:** 10.1101/471342

**Authors:** Ewart Kuijk, Francis Blokzijl, Myrthe Jager, Nicolle Besselink, Sander Boymans, Susana M. Chuva de Sousa Lopes, Ruben van Boxtel, Edwin Cuppen

**Author notes:** Equal contribution (Ewart Kuijk, Francis Blokzijl).

## Abstract

A developing human fetus needs to balance rapid cellular expansion with maintaining genomic stability. Here, we accurately quantified and characterized somatic mutation accumulation in fetal tissues by analyzing individual stem cells from human fetal liver and intestine. Fetal mutation rates were ~5-fold higher than in tissue-matched adult stem cells. The mutational landscape of fetal intestinal stem cells resembled that of adult intestinal stem cells, while the mutation spectrum of fetal liver stem cells is distinct from stem cells of the fetal intestine and the adult liver. Our analyses indicate that variation in mutational mechanisms, including oxidative stress and spontaneous deamination of methylated cytosines, contribute to the observed divergence in mutation accumulation patterns and drive genetic mosaicism in humans.

**One Sentence Summary:** Liver and intestinal cells accumulate elevated amounts and diverged types of somatic DNA mutations during early human fetal development

---

Mutations that arise during fetal development result in somatic mosaicism and can affect a large population of cells in the adult organism. Potential consequences for human health are congenital disorders and increased cancer risk (*1–4*). Understanding the processes that drive somatic mosaicism in development would therefore expand our knowledge on the origin and evolution of diseases. A powerful method for obtaining insight into active and past mutational processes is through the analysis of mutational profiles, which reflect balances between DNA damage and activity of DNA repair pathways. So far, 30 mutational signatures have been extracted from the mutational landscapes of cancer genomes and listed in the COSMIC database (*5*). Mutational signature analysis of human adult stem cells (SCs) has revealed that mutation accumulation in healthy cells is tissue-specific and happens with a continuous rate throughout adult life (*6*). Adult intestinal SCs have a large contribution of COSMIC Signature 1, which is characterized by C to T changes at CpG sites and has been attributed to spontaneous deamination of methylated cytosines. In contrast, adult liver SCs show minimal Signature 1 contribution, while Signature 5, characterized by T to C transitions, is prevalent (*6*). Signatures 1 and 5 were found to accumulate with age and act in a clock-like manner (*7*) and were also found to explain mutations that occur very early in development in 2-cell stage embryos (*8*). To date, it is unknown what the contribution of fetal development to cell-specific mutation accumulation is and when in life patterns of mutation accumulation start to diverge between tissues. Therefore, we accurately catalogued genome-wide *de novo* somatic mutations in individual liver and intestinal SCs of human fetuses at week 15-22 of gestation.

## Fetal mutation accumulation rates

Measuring extremely low mutation loads in individual cells is challenging, due to the high noise rates that are associated with single-cell DNA sequencing techniques that typically involve error-prone amplification methods (*9*). To circumvent this hurdle, we studied mutation accumulation in individual SC cells, by expanding them in vitro as organoids before whole genome sequencing and bioinformatic filtering for clonal variants - a method which we previously applied successfully to identify somatic variants in adult SCs from different tissues and donors of different ages (*6, 10*). Using routine organoid culture conditions (*11, 12*), we successfully derived SC lines from the fetal liver and intestine of four fetuses at weeks 15, 17, and 22 (n=2) of gestation. Individual organoids from the primary cultures were manually picked, expanded to obtain 14 clonal lines, 6 of the intestine and 8 of the liver (Fig. S1), and whole genome sequenced to a minimum of 30x average coverage. No chromosomal aberrations and aneuploidies were observed in the copy number profiles (data not shown). At the basepair level (see Materials and Methods for details), we identified a total of 834 somatic base substitutions in 14 SCs from 4 independent fetuses (Table S1). Independent validation of 569 base substitutions confirmed 556 (98%) of the variants as true positives (Table S2).

On average, each SC accumulated 67 base substitutions. Particularly for liver there was a high degree of variation (minimum = 20, maximum = 153), likely caused by the spread in fetal age, as there was little variation between fetuses of the same age (Fig. 1A). A linear mixed model analysis confirmed a significant correlation (corrected *P*-value = 0.04) between the number of base substitutions in the fetal liver and fetal age (Fig. 1A), indicating continuous accumulation of increasing numbers of mutations over time. Since such mutation accumulation rates SC have been measured previously for adult liver and intestine (*6*), we can now for the first time directly compare mutation accumulation rates between human adult and fetal SCs of the same organ. For 13 of the 14 fetal stem cells, the mutation rates fell outside the 95% confidence intervals of the slope estimates of the adult liver and small intestine SCs (linear mixed model, Table S1), with ~5-fold more somatic variants per week of fetal life than per week of adult life (Fig. 1B).

**Fig. 1.**
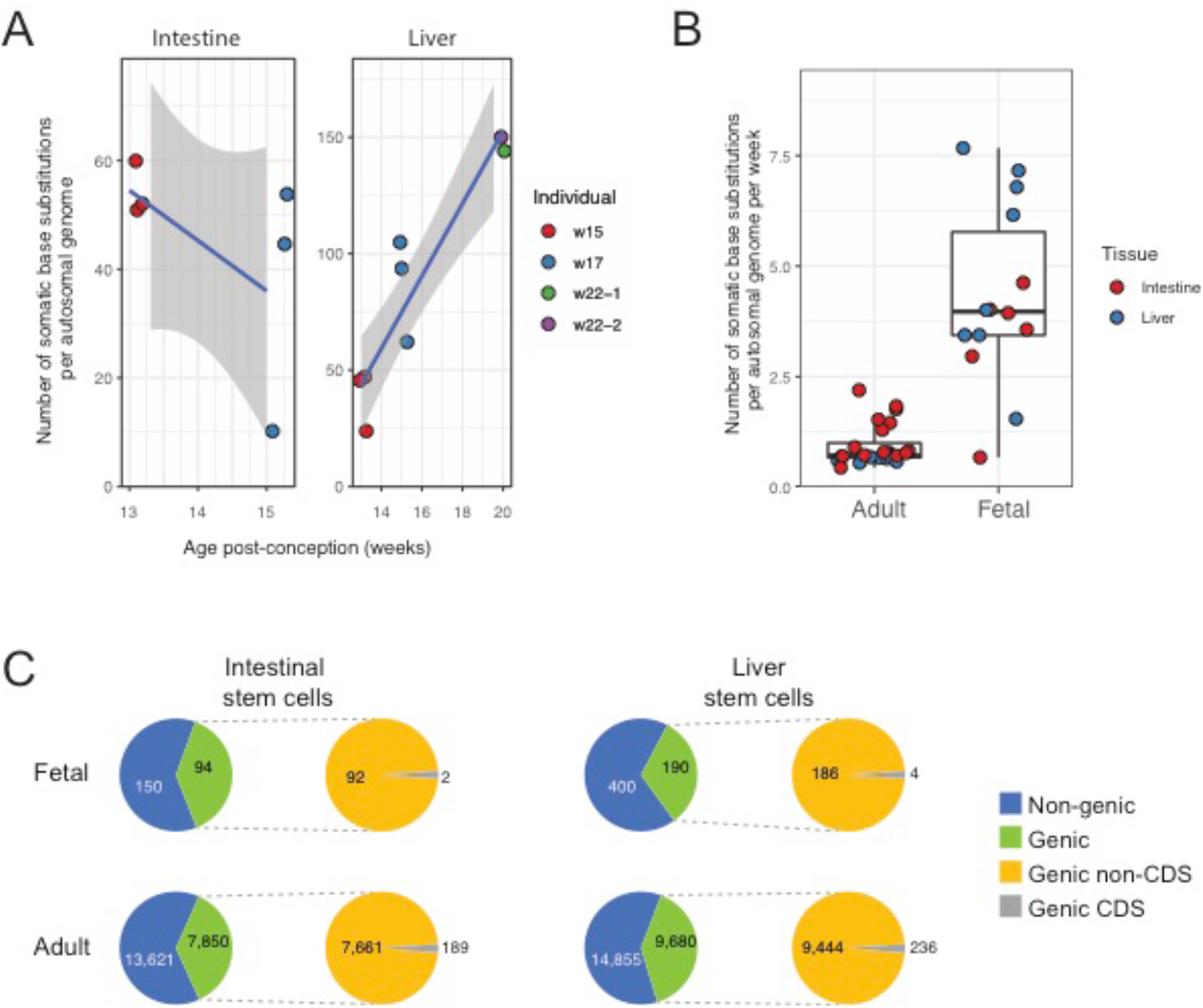
Mutation accumulation in stem cells of the human fetal liver and intestine. **(A)** Left panel: number of somatic base substitutions in each intestinal fetal stem cell (extrapolated to the whole autosomal genome). Right panel: number of somatic base substitutions in each liver fetal stem cell (extrapolated to the whole autosomal genome) as a function of fetal age in weeks postconception. Colors indicate the different fetuses. **(B)** Number of somatic base substitutions that accumulated per week in stem cells of adult and fetal liver and intestine. Each stem cell is represented by a data point. **(C)** Genomic location of somatic base substitutions for adults and fetuses and per indicated cell type. CDS = Coding sequence.

The observed mutation accumulation rates in SCs of the fetal liver and intestine (mean = 4) are well below the ~36 base substitutions per week that have been described for neural progenitors of similarly aged human fetuses (*13*). Thus, in contrast with what has recently been hypothesized (*13*), mutation rates in prenatal development appear not equal between tissues, although effects of technical differences in measurement methods cannot be excluded. Nevertheless, elevated mutation accumulation rates appear to be general for fetal cells of different tissues suggesting that the rapid cellular expansion in development comes at the cost of increased mutation accumulation. Unlike neural progenitors, SCs of the liver and the intestine fulfil important roles throughout adult life in tissue self-renewal and regeneration upon damage, thereby running a lifelong risk that somatic mutations are propagated to daughter SCs and contribute to tumor development. It may therefore be not surprising that these cells maintain higher levels of genomic integrity in early development.

The proportion of base substitutions in intergenic, genic, and protein-coding sequences is found to be similar for SCs from both fetal tissues, and comparable to the ratios found in adult SCs (Fig. 1C). The majority of the identified base substitutions are located outside genes. In fetal SCs, five nonsynonymous and one nonsense mutations were detected in protein coding sequences of six different genes (Fig. 1C, Table S3). None of these genes are involved in cell proliferation or have been causally implicated in cancer according to the COSMIC cancer gene census (*14*), suggesting that there was no positive selection for cells with functional somatic mutations that confer a selective advantage.

## Mutational patterns of fetal somatic mutations are tissue-specific

The mutation spectrum (Fig. 2, Fig. S2) of the fetal intestinal SCs closely resembled that of adult intestinal SCs (cosine similarity = 0.94) and was characterized by a large fraction of C to T changes, particularly at CpG sites and thus likely resulting from deamination of methylated cytosines. This similarity suggests that the balance in activity of DNA damage and repair processes is similar in adult and fetal intestinal SCs (Fig. 2B). In contrast, the spectrum of the fetal liver SCs was different between individuals of different ages (*P* < 0.001, Pearson’s Chi-squared test; Fig. 2A) and was distinct from that of adult liver SCs (*P* < 2.2e-16 Pearson’s Chi-squared test; Fig. 2A). Notably, the cosine similarity between the fetal liver and the fetal intestine was 0.72 (Fig. 2B), demonstrating that these tissues accumulate different types of mutations during fetal development and adult life. Since the liver mutation spectrum is variable with fetal age, we also compared all profiles separately. Although fewer base substitutions can be used per profile, which comes at the cost of reduced statistical power, comparison of the individual profiles confirmed the high resemblance between the fetal and adult intestinal SCs and the difference between the fetal liver and the fetal intestine (Fig. S3). The mutational profile of week 15 fetal liver SCs was most similar to the intestinal SCs, but this resemblance decreased with fetal age (Fig. 2C). Our results thus show that patterns of mutation accumulation between tissues already start to diverge prior to week 15 of gestation. This divergence seems to be caused by a shift in fetal liver SCs from C to T changes towards more C to A substitutions (Fig. 2A), similar as described for neural progenitors (*13*).

**Fig. 2.**
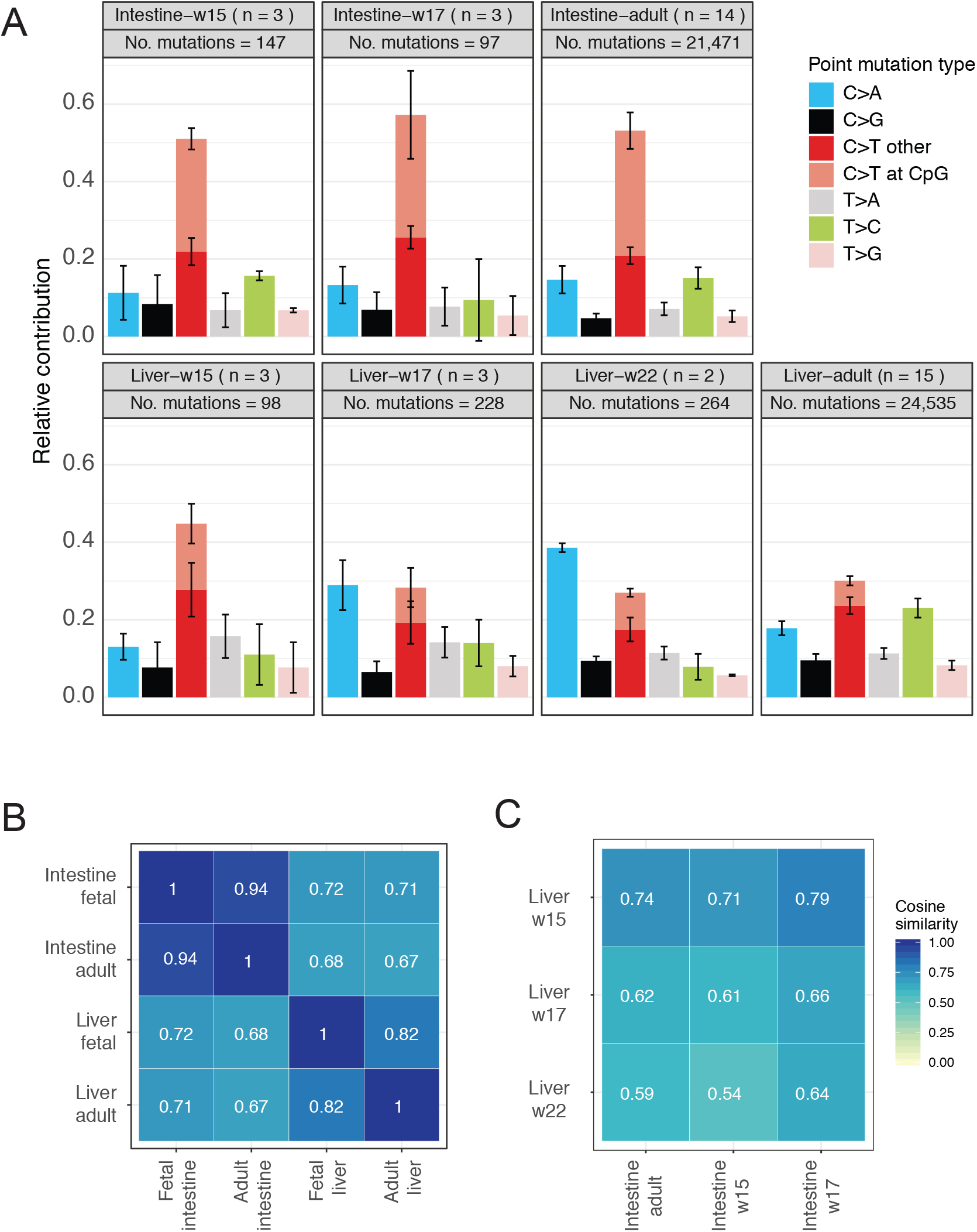
The fetal liver and fetal intestine have distinct mutational patterns. **(A)** Mutation spectra for all tissues and ages. Error bars represent standard deviations. The total number of identified somatic base substitutions per spectrum is indicated. **(B)** Cosine similarities between the average 96-type mutational profiles of liver and intestinal stem cells from fetal and adult origin. **(C)** Cosine similarities between the (average) 96-type mutational profiles of intestinal stem cells and liver stem cells of differently aged-fetuses.

## Fetal mutational Signature analysis

To gain further insight into the molecular processes underlying the mutation accumulation in fetal tissues, we compared the mutational profiles of the fetal SCs with the pan-cancer derived COSMIC mutational signatures (*15*). Unsupervised hierarchical clustering based on the cosine similarities of the mutational profiles of the samples and sample groups with the COSMIC signatures resulted in clustering of the SCs by tissue of origin. However, week 15 liver SCs are notable exceptions, as these are more similar to the intestinal samples (Fig. 3A). To expand on these findings, we reconstructed the mutational profiles of the adult and fetal SCs with the COSMIC signatures (*16*). The cosine similarities of the reconstructed profiles with the fetal SC profiles were 0.89-0.94 (Fig. S4), from which we conclude that the mutational spectrum can to a large extent be explained by the known COSMIC signatures. The residual part that is as yet unexplained might have a technical basis due to the low number of mutations, or could be caused by additional, unassessed mutational processes specific to fetal development (*16*).

**Fig. 3.**
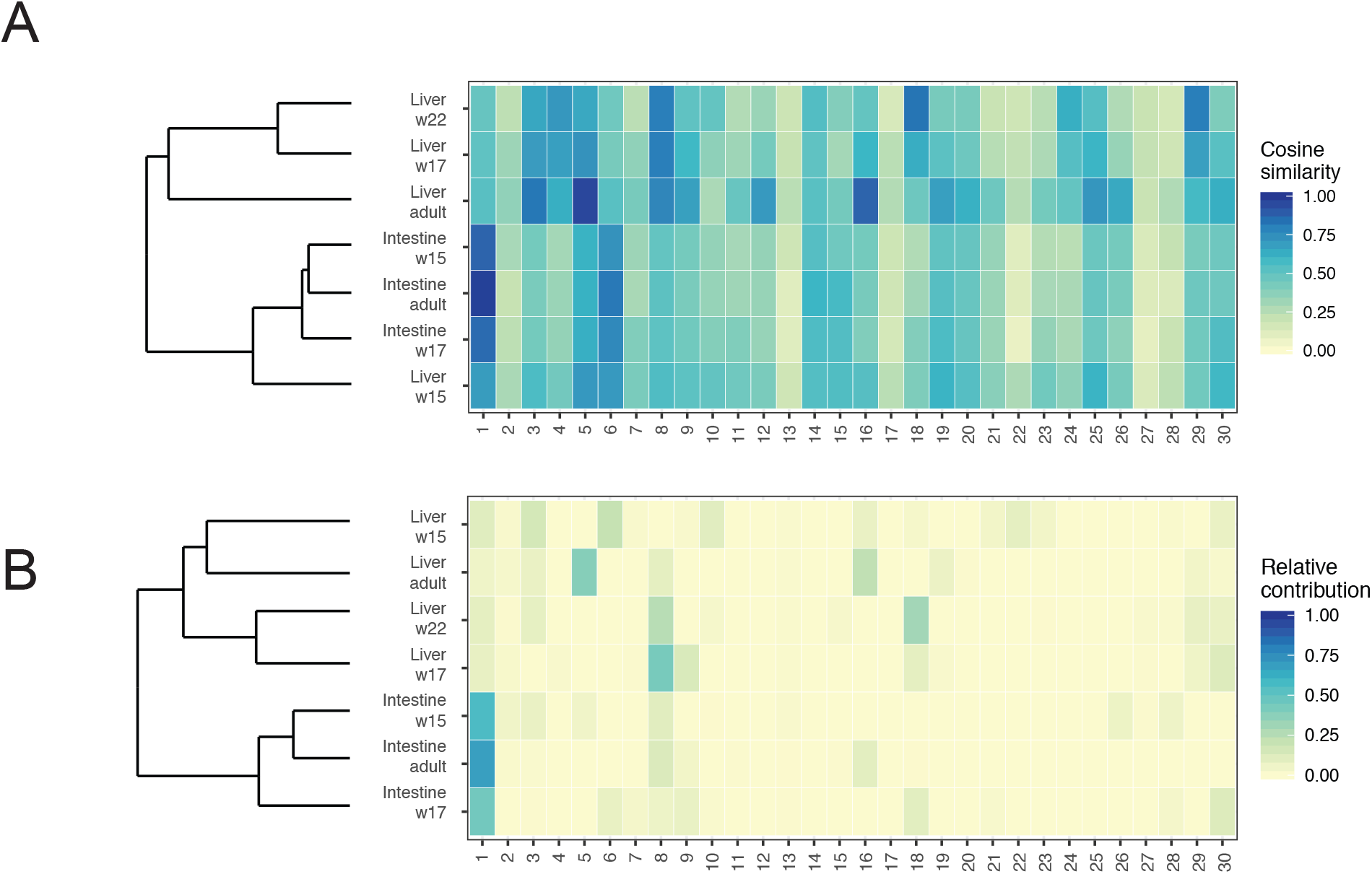
Resemblance between mutational profiles and COSMIC Signatures. **(A)** Cosine similarity heatmap between the COSMIC signatures and the mutational profiles of the adult and fetal stem cells. Samples are grouped by unsupervised hierarchical clustering. **(B)** Relative contribution heatmap of the COSMIC signatures to the mutational profiles of the adult and fetal stem cells. Samples are grouped by unsupervised hierarchical clustering.

The mutation profile of fetal, as well as adult intestinal SCs is most similar to COSMIC mutational Signature 1 (Fig. 3A). In contrast, there was only a minimal contribution of Signature 1 to the mutational profile of the rapidly proliferating fetal liver (*17*), particularly at weeks 17 and 22. This conflicts with the suggestion that cell types with a high cell division rate exhibit more Signature 1 mutations and that this Signature could serve as a clock that registers the number of mitoses that a cell has experienced since fertilization (*7*). Our results indicate that a high cell turnover rate is, at least in fetal development, not necessarily associated with a pronounced contribution of Signature 1-like mutations.

The fetal liver mutation profiles show resemblance to COSMIC Signature 8 and Signature 18, both of which are characterized by C to A changes, while adult liver SCs are highly similar to COSMIC signatures 5 and 16, the etiology of which is unknown (Fig 3). The etiology of Signatures 8 and 18 is also unknown, but has been linked to oxidative stress related mechanisms (*18*). These findings demonstrate that in fetal liver SCs other mutational mechanisms are dominant than in adult liver SCs.

To avoid biases originating from the existing COSMIC signatures, we performed a *de novo* extraction of mutational signatures using non-negative matrix factorization, using the mutational landscapes of all adult and fetal samples as input (*15*). We identified 3 signatures, of which Signature A turned out to be highly similar to COSMIC signatures 5 and 16, Signature B resembled COSMIC Signature 1, and Signature C showed resemblance to COSMIC Signatures 8 and 18 (Fig. S5). The extracted signatures confirm that the mutational profiles of fetal intestinal SCs resemble COSMIC Signature 1 and fetal liver SCs resemble COSMIC Signatures 8 and 18 (Fig. S5). At the individual sample level, the de novo signature analyses do confirm the conclusion that mutational profiles of fetal intestinal SCs bear resemblance to that of adult intestinal SCs, while fetal liver SCs have profiles distinct from adult liver, but also from fetal and adult intestine (Fig. S5B).

Since the fetal intestine and liver do not yet fulfill their adult tissue functions, the observed divergence between these tissues in mutation accumulation is probably not caused by differences in exposure to exogenous mutagens, but is more likely the result of cell-or tissue-intrinsic processes that are regulated differently between the developing organs. Principal component analysis of RNA-sequencing data revealed similar overall transcriptome profiles for fetal and adult intestinal SCs, whereas the fetal liver SCs are distinct from the intestinal SCs and the adult liver SCs (Fig. 4A). Unsupervised hierarchical clustering of the top-100 differentially expressed genes yielded similar results (Fig. 4B) and corresponded well with the clustering based on mutational signatures (Fig. 2B).

**Fig. 4.**
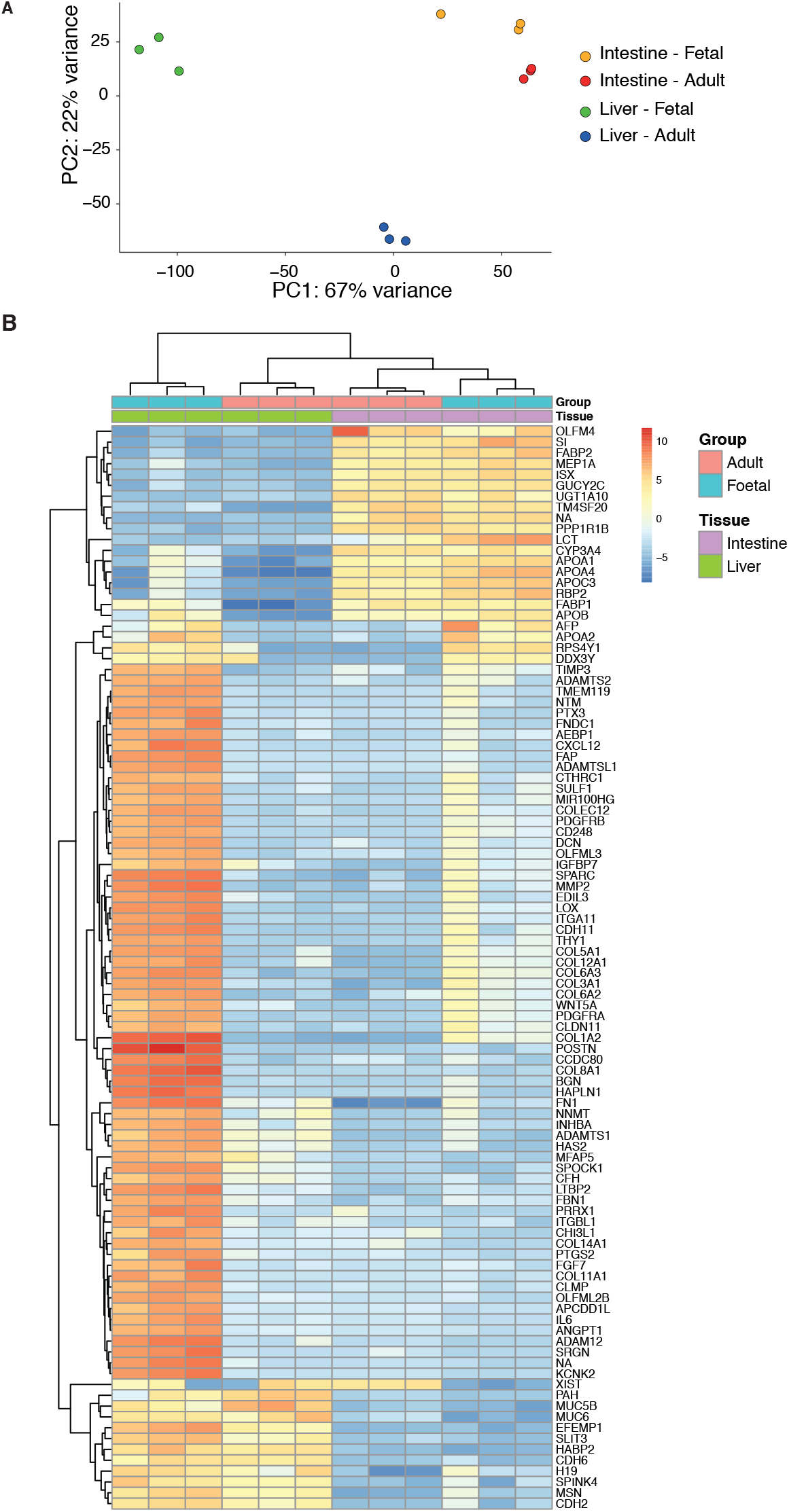
RNA-expression analysis. **(A)** Principal component analysis of the gene expression profiles. The individual samples are projected onto the first two principle components. PC1 = principal component 1, PC2 = principal component 2. **(B)** Gene expression heatmap of the top 100 differentially expressed genes between adult and fetal liver and intestinal stem cells. Genes are ordered by non-supervised hierarchical clustering.

To gain more insight into putative mechanisms leading to fetal tissue-specific mutation accumulation, we focused on the dominant observed substitution types, i.e. C to A changes in the fetal liver and C to T changes at CpG sites in the fetal intestine. C to A transversions can result from DNA damage by oxidation, resulting in 8-oxoguanine pairing aberrantly with adenine (*19*). *MUTYH* and *OGG1* are the major glycosylases that identify oxidized bases and initiate base excision repair (BER) (*20*). Inactivating mutations in these genes lead to more C to A substitutions (*21, 22*). The expression of *MUTYH* and *OGG1* is lower in the fetal liver than in the fetal intestine (Fig. S6), which could result in increased C to A transversions in the fetal liver.

The predominant base changes in fetal and adult intestinal SCs (C to T changes at CpG sites) are also frequent in early embryogenesis (*8, 23*). The underlying cause, spontaneous deamination of methylated cytosines, therefore appears to be a dominant mutagenic factor from fertilization until adulthood in intestinal SCs and their progenitors. Spontaneous deamination of methylated cytosines at CpG dinucleotides results in T:G mismatches (*5*). These mismatches can be effectively repaired by BER (*24*) and potentially non-canonical mismatch repair (MMR)(*25*), but a mutation becomes fixed if the cell divides before the mismatch is resolved, resulting in a C to T change (*7*). Methyl-CpG binding domain 4 (MBD4) and Thymine DNA Glycosylase (TDG) are the two major glycosylases that specifically catalyze the removal of thymines opposite of guanines at CpG sites (*24, 26*). Both glycosylases are expressed at higher levels in SCs of the fetal liver than in SCs of the fetal intestine (Fig. S6). This may lead to tissue specific rates of thymine excision at the T:G mismatches by BER resulting in differences in C to T substitution rates at CpG sites. These results suggest that tissue-specific mutation accumulation in fetal development may indeed be the result of cell-intrinsic processes such as differential expression of BER glycosylases, although the distinction between cause and consequences remains difficult and would require induction and measurement of DNA damage and the activity of DNA repair in fetal tissues.

In conclusion, our results show that fetal growth comes at the cost of elevated mutation rates. Furthermore, distinct mutational mechanisms shape the mutational landscapes of the fetal liver and the fetal intestine. Spontaneous deamination of methylated cytosines is the main driver of mutation accumulation in fetal intestinal stem cells, while oxidative mechanisms are important for mutation accumulation in fetal liver stem cells.

## Acknowledgments

The authors thank the Centre for Contraception, Abortion and Sexuality (CASA) in Leiden and the Hague for their efforts to obtain the human fetal material and the USEQ for their support with next-generation sequencing experiments; the authors would also like to thank Luc J. W. van der Laan of the Erasmus MC for providing adult liver samples and Sabine Middendorp for providing adult intestinal samples, for which previously published data has been used in the current study.

## Author contributions

E.K., M.J. and N.B. performed wet-lab experiments, E.K., F.B., M.J. and S.B. performed bioinformatic analyses. S.M.C.d.S.L. obtained and isolated fetal tissues. E.K., F.B., M.J., R.v.B. and E.C. were involved in the conceptual design of the study. E.K., F.B., M.J. and E.C. wrote the manuscript;

## Data and materials availability

The RNA and DNA sequencing data have been deposited at the European Genome-Phenome Archive (EGA) under accession numbers EGAS00001002886, EGAS00001001682, and EGAS00001002983.

## Supplementary materials

Materials and Methods Figures S1–S6 Tables S1–S3

## Supplementary Materials

### Materials and Methods

#### Collection of human material

The Medical Ethical Committee of the Leiden University Medical Center approved this study (P08.087). Informed consent was obtained on the basis of the Declaration of Helsinki (World Medical Association). Human fetal tissues (intestine, liver and skin) at gestational ages W15-W22 were collected from elective abortion material (vacuum aspiration) without medical indication. In this study, weeks of gestation, which is equivalent to the last menstrual period (LMP), was determined by utrasonography. The age in weeks post-conception was determined by subtracting two weeks from the weeks of gestation. After collection, the material was washed with 0.9% NaCl (Fresenius Kabi, France) and stored on ice until further processing.

#### Fetal stem cell isolation and clonal organoid culture

Organoid cultures from liver and intestinal tissue material were derived as previously described (*1, 2*). In short, liver biopsies were minced and subsequently dissociated into single cell solutions using human liver digestion solution (EBSS supplemented with 1 mg/ml collagenase type 1A and 0.1 mg/ml DNaseI). Cells were plated at limiting dilution and organoid cultures were initiated by culturing these cells in BME overlaid with human liver isolation medium (60% Advanced DMEM/F-12 (supplemented with 1% penicillin/streptomycin, 1% GlutaMAX, and HEPES 10 mM), 30% WNT3A conditioned medium (produced in house), 10% RSPOI conditioned medium (produced in house), 1x B27 supplement without retinoic acid, 1x N2 supplement, 1x Primocin, 1 : 1,000 hES cell cloning & recovery supplement, 10 mM Nicotinamide, 1.25 mM N-acetylcysteine, 5 μM A83-01, 10 μM Forskolin, 10 μM Rho kinase inhibitor, 10 nM Gastrin I, 100 ng/ml Noggin, 100 ng/ml FGF10, 50 ng/ml hEGF, and 25 ng/ml HGF). For the intestine, villi were removed from intestinal biopsies by applying mechanical strength. Subsequently, the biopsies were minced and dissociated into single cell solutions by incubating these tissue pieces with Tryple at 37°C for 10-20 minutes. Cells were plated at limiting dilution and organoid cultures were initiated by culturing these cells in matrigel overlaid with human intestinal isolation medium (30% Advanced DMEM/F-12 (supplemented with 1% penicillin/streptomycin, 1% GlutaMAX, and HEPES 10 mM), 50% WNT3A conditioned medium (produced in house), 20% RSPOI conditioned medium (produced in house), 1x B27 supplement, 1x Primocin, 1 : 1,000 hES cell cloning & recovery supplement,10μM SB 202190, 10 mM Nicotinamide, 1.25 mM N-acetylcysteine, 0.5 μM A83-01, 10 μM Rho kinase inhibitor, 100 ng/ml Noggin, and 50 ng/ml hEGF). After 2-3 days, organoids started to appear and the medium was changed to human liver expansion medium and human intestinal expansion medium, respectively (recipe as described above, without Rho kinase inhibitor and hES cell cloning & recovery supplement). Clonal cultures were derived by picking single organoids and expanding them until there was sufficient material to perform WGS.

#### Whole genome sequencing and variant identification

DNA was isolated from the organoid cultures and skin and liver tissue using the QIAsymphony DNA mini kit (Qiagen). DNA libraries for Illumina sequencing were generated from 200 ng genomic DNA using standard protocols (Illumina) and sequenced 2 x 100 bp paired-end to 30X base coverage with the Illumina HiSeq Xten at the Hartwig Medical Foundation. WGS data was processed and copy number analyses and base substitutions were called using GATK (*3*) and filtered as described previously (*4*). Skin was used as a control in the filtering pipeline. Because for one skin sample, not enough DNA could be retrieved, a bulk tissue liver sample was used as reference instead. In addition to the filtering described in (*4*), we filtered on genotype quality (GQ): GQ >=99 in the clonal organoid culture, and GQ >=10 in the polyclonal control sample, as these positions commonly represent FP somatic variants. To generate a set of high confident variants, we visually inspected all base substitutions using the Integrative Genomics Viewer (IGV) (*5*) and excluded 44% of all base substitutions. Base substitutions in repetitive sequences (e.g. poly-G stretches), nearby an indel (within ~20bp of base substitution), in regions with a very high coverage (as these are most likely mismapped reads), that were found in the majority of the other clones (as these are most likely missed germline positions), and within a deletion or duplication were flagged as false positives. For adult SCs, we used data from previously established human liver and intestinal stem cell lines (*6, 7*).

For 569 of the variants that were identified in fetal SCs, 1,138 custom capture probes of 120 bp were designed (one for the variant and one for the reference allele) and ordered. DNA sequencing libraries were prepared according to the manufacturer’s (Twist Bioscience) protocol. In short, DNA was enzymatically fragmented followed by end repair and dA-tailing. Subsequently, index adapters were ligated and a pre-capture PCR amplification was performed. Libraries were pooled and a bead-based size selection was performed. The size-selected libraries were hybridized to the capture probes. After binding to streptavidin beads, the enriched libraries were PCR amplified, purified and sequenced 2×150 bp on the NextSeq500. All sample were thus analyzed for all target loci, thereby functioning as each others controls. Variants were called using GATK (*3*) and considered positive if unique to the sample in which the sample was originally called. All other variants were manually inspected in IGV. This validation resulted in the confirmation of 98% of all variants (555 after initial automated filtering, 1 after manual inspection in IGV; Table S2). For the 13 remaining variants the results were inconclusive, because the custom capture failed at these positions.

#### Base substitution load and type

True positive base substitutions were extracted from the VCF files. The base substitution load was computed by extrapolating the number of mutations to the autosomal genome, as described previously (*4*). To allow comparison between the accumulation during fetal development and adult life, we downloaded VCF files and surveyed bed files of healthy liver and small intestine adult SCs from https://wgs11.op.umcutrecht.nl/mutational_patterns_ASCs/. In addition, mutation catalogues of five clonal liver organoid cultures derived from four different healthy liver donors were included in this study (Jager et al. personal communication; data available upon request).

Using the Linear and Nonlinear Mixed Effects Models R package version 3.1-137, the slope and 95% confidence intervals for the mutation rates of the adult liver and small intestine were determined. Next, we tested if the mutation rates per week post-conception fell within the 95% confidence interval.

Mutational pattern analysis was performed as described previously, using the MutationalPatterns R package (*8*). 7-type mutation spectra were extracted from the VCF files. 96-type mutational profiles were subsequently generated and the average profile (centroid) was determined for the assessed SC types. Centroids were reconstructed with the 30 COSMIC mutational signatures. Subsequently, signatures that contributed at least 5% to one or more centroids were selected and the centroids were reconstructed again using this subset of the mutational signatures. Cosine similarities between samples and/or signatures and relative contributions of signatures were calculated using MutationalPatterns (*8*).

Gene definitions for hg19 were retrieved from the University of California, Santa Cruz (UCSC) Genome Browser (*9*) as a TxDb annotation package from Bioconductor. Functional consequences of coding mutations were identified using the VariantAnnotation R package (*10*).

#### RNA sequencing

Organoid cultures at passage 5 were lysed with Trizol (Life Technologies) and RNA was isolated using the QIAsymphony RNA kit (Qiagen). mRNA was isolated from 50 ng total RNA and libraries were subsequently generated using the Illumina Neoprep TruSeq stranded mRNA library prep kit (Illumina, NP-202-1001). mRNA libraries were sequenced 1 x 75 bp on the NextSeq500.

RNA sequencing reads were mapped with STAR v.2.4.2a to the human genome assembly GRCh37. The BAM files were sorted with Sambamba v0.5.8 and reads were counted with HTSeq-count version 0.6.1p1 (default settings) to exons. DESeq2 v_1.16.1 was used to normalize counts and for rlog transformation, followed by non-supervised hierarchical clustering of the top 100 differentially expressed genes (*11*). Principal component analysis was performed on normalized and rlog transformed counts. Gene ontology enrichment analysis was performed using the clusterProfiler package version 3.8.1 (*12*).

---

**Fig. S1.**
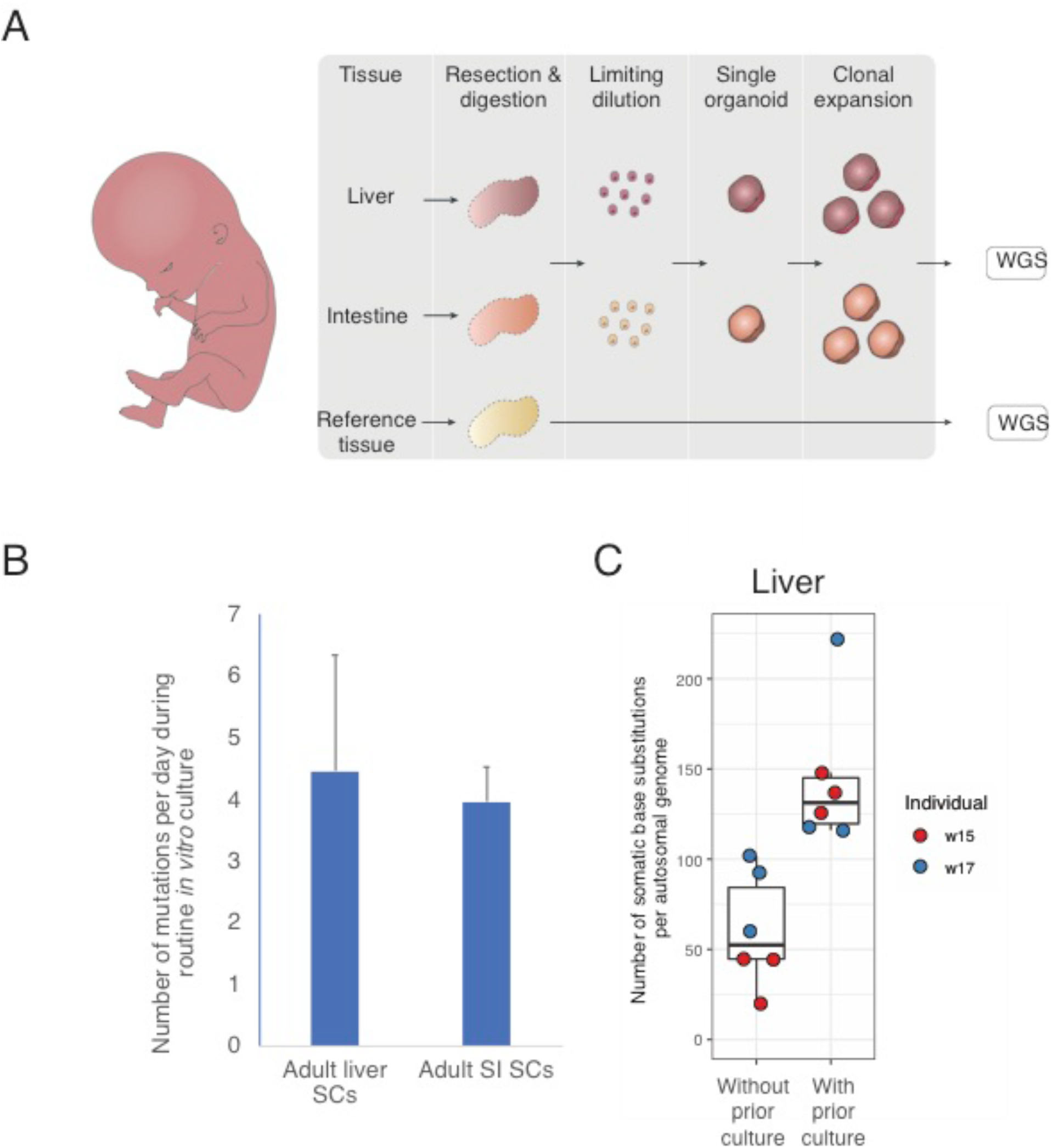
Experimental set-up. **(A)** Fetal liver and intestinal tissue samples were minced and enzymatically digested to obtain a single-cell suspension that was plated according to a limiting dilution series. Individual organoids were picked from the lowest concentrations, clonally expanded and subjected to whole genome sequencing. A multilineage bulk reference tissue from the same fetus was also sequenced to identify all germline variants, which were subsequently subtracted from all the variants in the stem cell clone. This protocol is slightly different from our previous studies (*4, 6*), in which we shortly cultured the cells (7-14 days) between isolation and the clonal step. Culturing before the clonal step was avoided, since culture-induced mutations would likely overshadow the very few mutations acquired *in vivo*. A minor disadvantage to this modified protocol is that more than one cell contributed to the individually expanded organoids. This could lead to a slight underestimation of the total number of mutations, because we would be looking at base substitutions of the *in vivo* common ancestor. A major advantage is that all measured mutations have originated *in vivo* and are not *in vitro* artefacts, thereby enabling accurate mutational profiling. (B) Adult liver and intestinal SCs acquire on average 4 additional base substitutions per day *in vitro*, which was avoided in the fetal SCs by generating clonal organoid cultures without prior *in vitro* culture. To determine *in vitro* mutation accumulation rates, clonal liver stem cells (n=4) and intestinal stem cells (n=6) were cultured for ~3 months after which a second clonal step was performed. Clones, subclones and bulk samples were whole genome sequenced and the number of mutations in between the clonal steps was determined by subtracting all variants in the clone and the bulk from the subclone. Error bars represent standard deviation. (C) Fetal liver stem cells that were cultured prior to the clonal step acquire more mutations than fetal liver stem cells for which the clonal step was performed immediately after isolation. This again demonstrates the necessity to avoid *in vitro* culture if the number of mutations is extremely low, as is the case for fetal cells. Of note, adult stem cells have many more mutations, and therefore the effect of a short period of *in vitro* culture is neglectable.

**Fig. S2.**
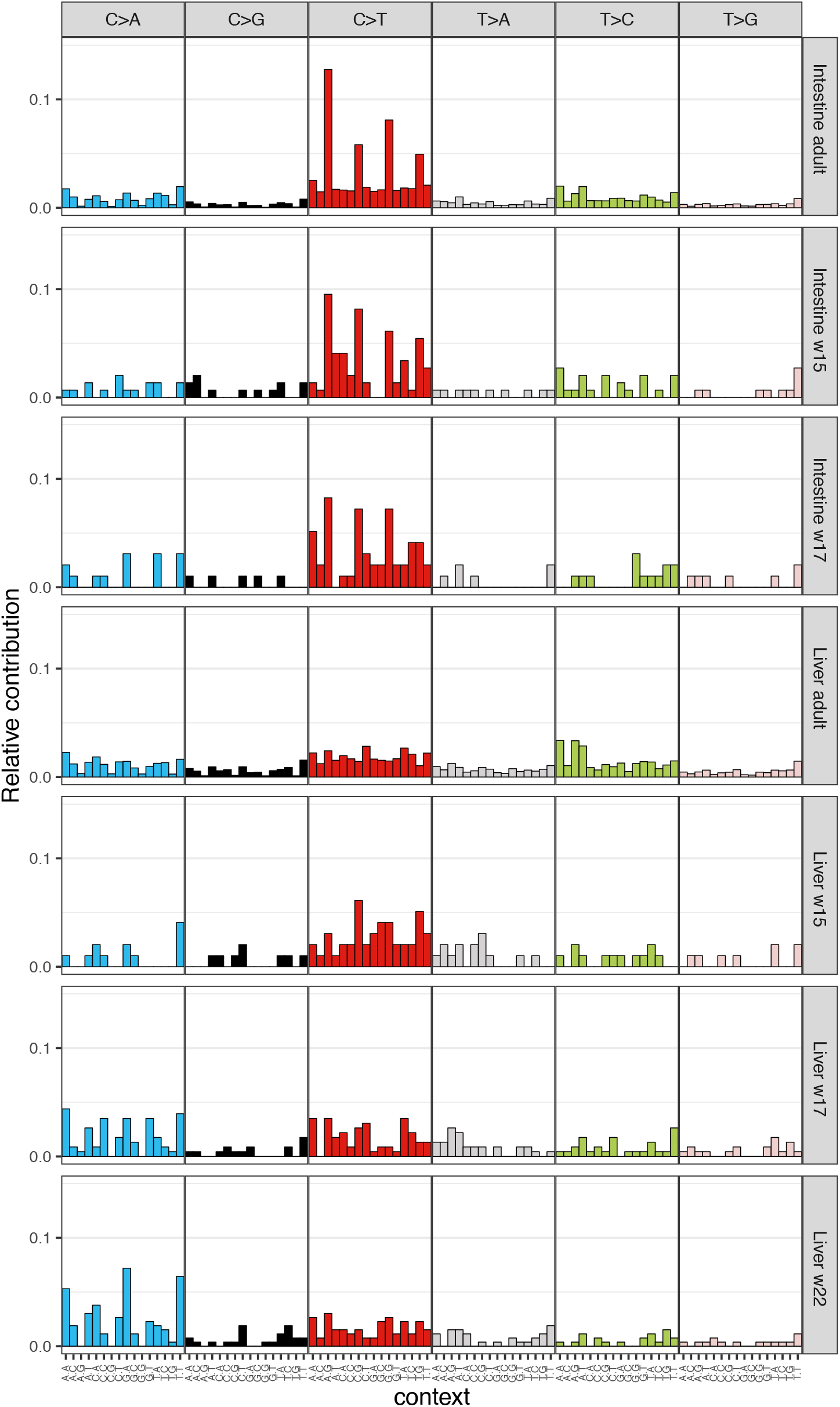
Mutational patterns for each tissue and age

**Fig. S3.**
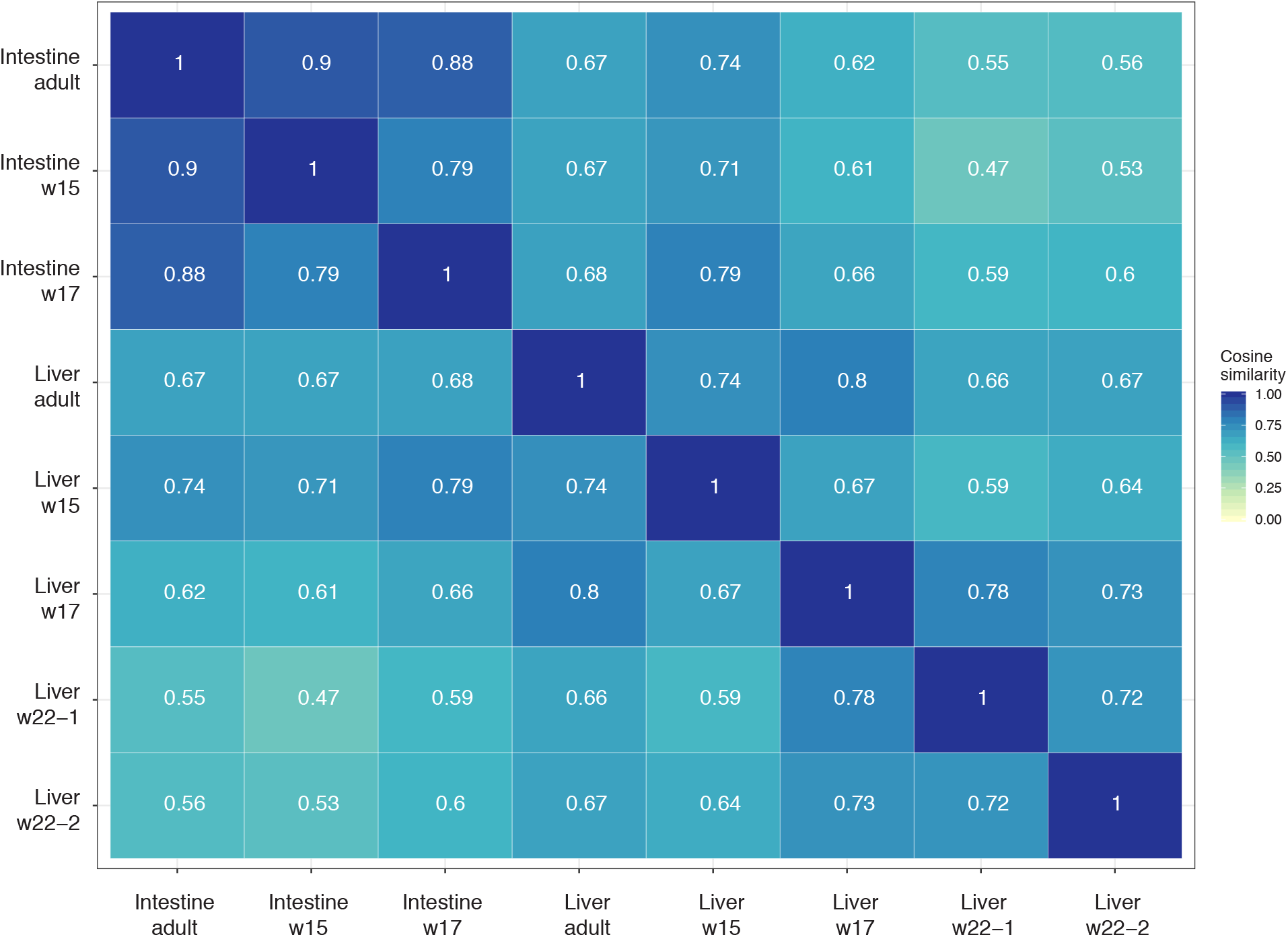
Cosine similarity heatmap. A heatmap showing the cosine similarity between the mutational profiles of all the indicated samples.

**Fig. S4.**
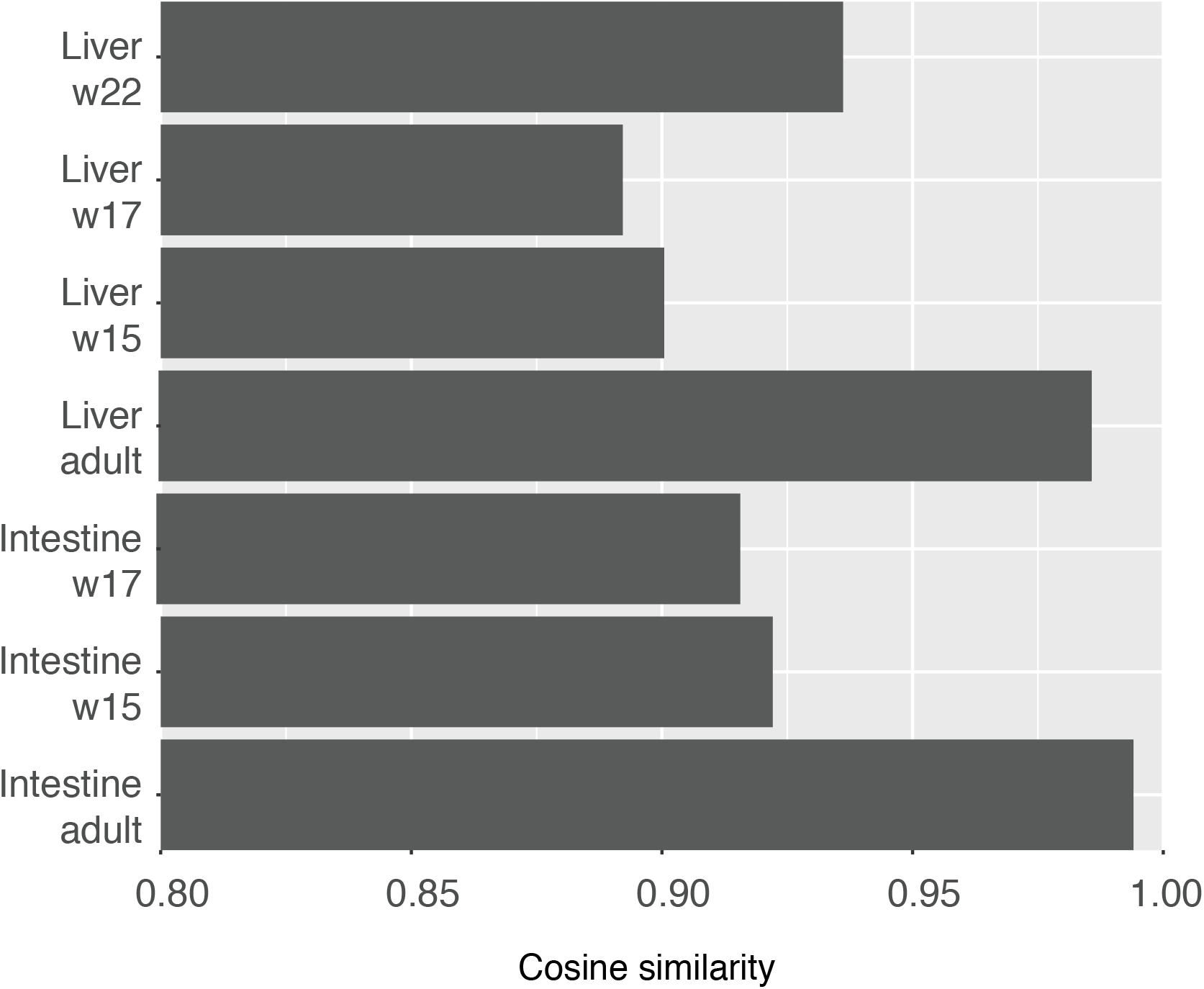
Reconstruction— of mutational profiles using known mutational signatures. The cosine similarity between the original mutational profile and the reconstructed mutational profile based on the optimal linear combination of all 30 COSMIC signatures.

**Fig. S5.**
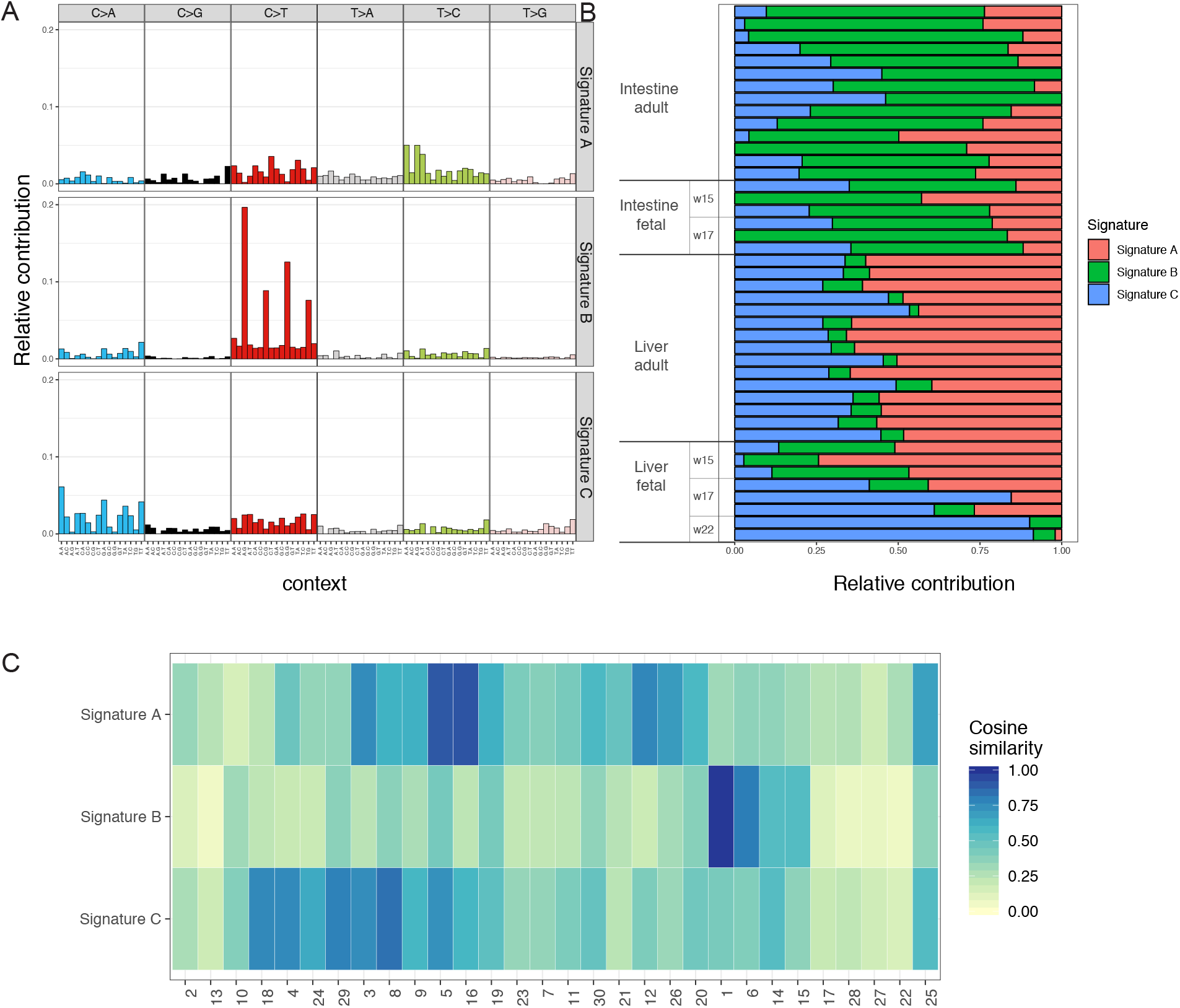
*De novo* identification of mutational signatures. **(A)** Mutational profiles of three signatures, extracted from the mutational profiles of all fetal and adult SCs using non-negative matrix factorization. The three extracted signatures are similar to the signatures that were previously described for adult liver and intestinal SCs (*6*). Signature A is characterized by T to C changes. Signature B is characterized by C to T changes at CpG sites. Signature C is characterized by C to A changes. **(B)** The relative contribution of the three signatures depicted in (A) to all individual SCs that have been analyzed in the current study. Signature A is predominantly found in adult liver SCs. Signature B is predominantly found in adult and fetal intestinal SCs, but also contributes to week 15 fetal liver samples. Signature C contributes to the mutational profile of fetal liver SCs from week 17 and week 22. (C) Cosine similarity heatmap between the *de novo* extracted signatures depicted in (A) and all 30 signatures from the COSMIC database. The order of the COSMIC Signatures is based on unsupervised hierarchical clustering to group signatures by similarity. Signature A is reminiscent of COSMIC Signatures 5 and 16, Signature B resembles COSMIC Signature 1, and Signature C is similar to COSMIC Signatures 8 and 18.

**Fig. S6.**
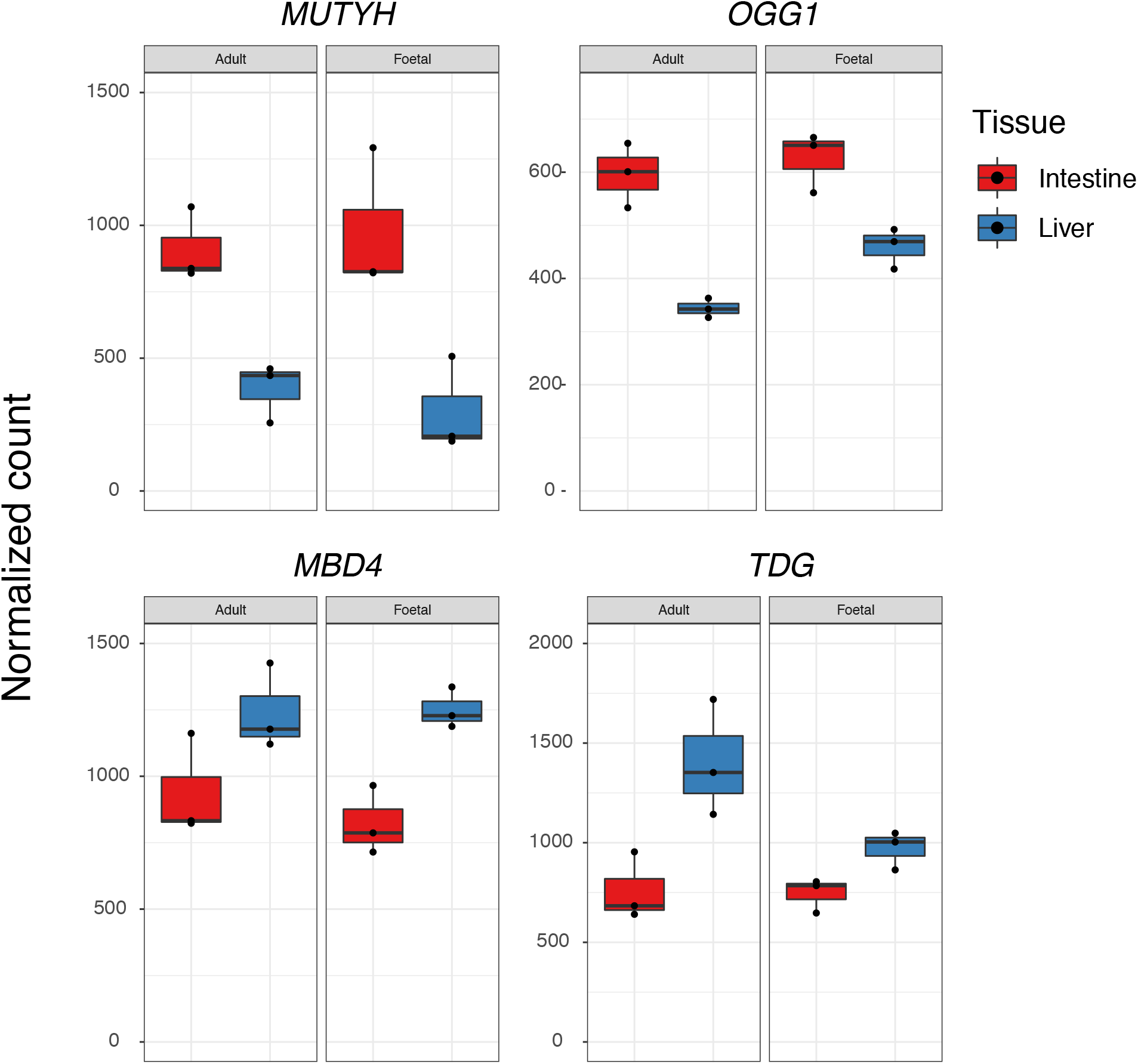
Gene expression of important DNA repair components. Normalized read counts for glycosylases involved in the repair of oxidized bases (MUTYH and OGG1) and in repair of T:G mismatches (MDB4 and TDG).

**Table S1.** Summary of all samples and the number of mutations identified per sample. Percentage surveyed based on an autosomal length of 2,881,033,286 bp.

| Individual | Tissue | Somatic variants | Surveyed (%) | Somatic variants per genome | weeks post conception | Somatic variants per genome per week | Within 95% CI of adult SCs |
| --- | --- | --- | --- | --- | --- | --- | --- |
| w22-1 | Liver | 127 | 88.71 | 143.2 | 20 | 7.16 | FALSE |
| w22-2 | Liver | 137 | 89.28 | 153.4 | 20 | 7.67 | FALSE |
| w15 | Intestine | 46 | 89.92 | 51.2 | 13 | 3.94 | FALSE |
| w15 | Intestine | 54 | 89.92 | 60.1 | 13 | 4.62 | FALSE |
| w15 | Intestine | 47 | 89.98 | 52.2 | 13 | 4.02 | FALSE |
| w15 | Liver | 40 | 89.48 | 44.7 | 13 | 3.44 | FALSE |
| w15 | Liver | 18 | 89.77 | 20.1 | 13 | 1.55 | FALSE |
| w15 | Liver | 40 | 89.44 | 44.7 | 13 | 3.44 | FALSE |
| w17 | Intestine | 48 | 89.7 | 53.5 | 15 | 3.57 | FALSE |
| w17 | Intestine | 9 | 89.55 | 10.1 | 15 | 0.67 | TRUE |
| w17 | Intestine | 40 | 90.15 | 44.4 | 15 | 2.96 | FALSE |
| w17 | Intestine | 54 | 89.88 | 60.1 | 15 | 4.01 | FALSE |
| w17 | Intestine | 91 | 89.43 | 101.8 | 15 | 6.79 | FALSE |
| w17 | Intestine | 83 | 89.82 | 92.4 | 15 | 6.16 | FALSE |

**Table S2.** Results for the validation of 569 base substitutions by capture using custom designed probes, followed by resequencing.

| Chromosome | position | Validated | REF | ALT |
| --- | --- | --- | --- | --- |
| 1 | 102086547 | Yes | C | T |
| 1 | 102739619 | Yes | A | T |
| 1 | 104541988 | Yes | G | A |
| 1 | 104989439 | Yes | G | A |
| 1 | 106950585 | Yes | A | G |
| 1 | 108903606 | Yes | C | T |
| 1 | 110669415 | Yes | G | A |
| 1 | 113744543 | Yes | C | T |
| 1 | 118275369 | Yes | A | C |
| 1 | 154536891 | Yes | T | A |
| 1 | 162445971 | Yes | G | A |
| 1 | 163297623 | Yes | T | G |
| 1 | 164131242 | Yes | G | A |
| 1 | 167291858 | Yes | G | A |
| 1 | 170366385 | Yes | A | T |
| 1 | 171824998 | Yes | G | T |
| 1 | 173011638 | Yes | A | C |
| 1 | 174047834 | Yes | A | T |
| 1 | 174150412 | Yes | G | T |
| 1 | 17429855 | Yes | G | A |
| 1 | 18561684 | Yes | G | A |
| 1 | 185700548 | Yes | C | T |
| 1 | 186980520 | Yes | T | C |
| 1 | 188308049 | Yes | C | T |
| 1 | 19385285 | Yes | G | A |
| 1 | 196440886 | Yes | C | T |
| 1 | 198008740 | Yes | G | A |
| 1 | 212415996 | Yes | C | T |
| 1 | 214656654 | Yes | T | G |
| 1 | 219561897 | Yes | C | T |
| 1 | 229613599 | Yes | C | T |
| 1 | 241691591 | Yes | C | T |
| 1 | 242754428 | Yes | G | C |
| 1 | 27210699 | Yes | A | G |
| 1 | 30671653 | Yes | C | A |
| 1 | 31539814 | No | #N/A | #N/A |
| 1 | 33298575 | Yes | G | A |
| 1 | 37011967 | Yes | A | G |
| 1 | 44653231 | Yes | C | T |
| 1 | 60780422 | Yes | C | T |
| 1 | 61537077 | Yes | C | T |
| 1 | 70022130 | Yes | A | C |
| 1 | 72519523 | Yes | T | A |
| 1 | 75155678 | Yes | G | C |
| 1 | 81393714 | Yes | A | G |
| 1 | 82777183 | No | #N/A | #N/A |
| 1 | 89100411 | Yes | C | A |
| 1 | 97868091 | Yes | T | C |
| 10 | 102507556 | Yes | T | G |
| 10 | 104718210 | No | #N/A | #N/A |
| 10 | 107349738 | Yes | C | T |
| 10 | 107632620 | Yes | C | A |
| 10 | 108426308 | Yes | C | G |
| 10 | 117418886 | Yes | G | A |
| 10 | 119505937 | Yes | C | T |
| 10 | 120958767 | Yes | C | T |
| 10 | 125255937 | Yes | C | A |
| 10 | 126991664 | Yes | A | T |
| 10 | 132010654 | Yes | C | T |
| 10 | 2351000 | Yes | C | T |
| 10 | 24584983 | Yes | G | A |
| 10 | 30431190 | Yes | C | T |
| 10 | 58821708 | Yes | G | T |
| 10 | 62855802 | Yes | C | T |
| 10 | 64740294 | Yes | A | G |
| 10 | 71720520 | Yes | C | A |
| 10 | 78210363 | Yes | C | T |
| 10 | 81980307 | No | #N/A | #N/A |
| 10 | 81980308 | No | #N/A | #N/A |
| 10 | 8276500 | Yes | C | A |
| 10 | 85911581 | Yes | G | T |
| 10 | 87921276 | Yes | G | T |
| 10 | 9513321 | Yes | A | T |
| 10 | 95447789 | Yes | T | G |
| 11 | 100205843 | Yes | C | T |
| 11 | 111167109 | Yes | T | C |
| 11 | 119388191 | Yes | G | A |
| 11 | 123499990 | Yes | A | G |
| 11 | 16439179 | Yes | A | T |
| 11 | 17931809 | Yes | G | T |
| 11 | 1813085 | Yes | C | T |
| 11 | 28419268 | Yes | C | G |
| 11 | 30927611 | Yes | T | G |
| 11 | 37843396 | Yes | C | A |
| 11 | 38984046 | Yes | G | T |
| 11 | 48545587 | Yes | C | T |
| 11 | 48680274 | Yes | G | A |
| 11 | 50274133 | Yes | G | A |
| 11 | 50439820 | Yes | A | T |
| 11 | 5415787 | Yes | G | A |
| 11 | 56773350 | Yes | C | A |
| 11 | 57973572 | Yes | T | C |
| 11 | 59205380 | Yes | C | T |
| 11 | 6486329 | Yes | A | C |
| 11 | 65404068 | Yes | G | A |
| 11 | 80116117 | Yes | T | C |
| 11 | 835788 | No | #N/A | #N/A |
| 11 | 88535468 | Yes | A | C |
| 11 | 91443008 | Yes | G | A |
| 11 | 91468728 | Yes | C | A |
| 11 | 94108344 | Yes | G | C |
| 12 | 101207165 | Yes | G | T |
| 12 | 102630374 | Yes | C | T |
| 12 | 10452573 | Yes | C | T |
| 12 | 104844559 | Yes | T | G |
| 12 | 113717918 | Yes | C | T |
| 12 | 127708815 | Yes | G | A |
| 12 | 128653269 | Yes | C | T |
| 12 | 133063387 | Yes | C | T |
| 12 | 1412611 | Yes | G | T |
| 12 | 1481683 | Yes | G | A |
| 12 | 20882003 | Yes | C | T |
| 12 | 22026776 | Yes | T | A |
| 12 | 31870827 | Yes | C | G |
| 12 | 33309544 | Yes | C | A |
| 12 | 34790612 | Yes | G | T |
| 12 | 38096050 | Yes | G | A |
| 12 | 45002653 | Yes | G | T |
| 12 | 47036834 | Yes | T | C |
| 12 | 47036835 | Yes | A | C |
| 12 | 47665312 | Yes | G | A |
| 12 | 53814257 | Yes | T | G |
| 12 | 60191210 | Yes | G | A |
| 12 | 70749928 | Yes | T | A |
| 12 | 72944702 | Yes | G | T |
| 12 | 80491045 | Yes | G | A |
| 12 | 82732819 | Yes | A | G |
| 12 | 84111843 | Yes | C | A |
| 12 | 90339938 | Yes | C | T |
| 12 | 9087600 | Yes | G | A |
| 12 | 91476283 | Yes | C | T |
| 12 | 914979 | Yes | C | T |
| 12 | 94872655 | Yes | A | C |
| 13 | 112049895 | Yes | C | T |
| 13 | 19107026 | No | #N/A | #N/A |
| 13 | 21301355 | Yes | C | T |
| 13 | 24168730 | Yes | G | T |
| 13 | 26514296 | Yes | G | T |
| 13 | 28895836 | Yes | G | A |
| 13 | 43091864 | Yes | C | G |
| 13 | 53998452 | Yes | A | G |
| 13 | 56195367 | Yes | A | T |
| 13 | 58600506 | Yes | C | A |
| 13 | 64820987 | Yes | G | A |
| 13 | 65594528 | Yes | T | A |
| 13 | 67781266 | Yes | A | G |
| 13 | 69868826 | Yes | C | A |
| 13 | 70955590 | Yes | T | G |
| 13 | 72388403 | Yes | G | C |
| 13 | 72822156 | Yes | C | T |
| 13 | 73753835 | Yes | C | T |
| 13 | 74390356 | Yes | C | A |
| 13 | 77925561 | Yes | G | T |
| 13 | 83596121 | Yes | T | A |
| 13 | 84029220 | Yes | T | C |
| 13 | 84856486 | Yes | C | A |
| 13 | 86673916 | Yes | C | A |
| 13 | 86994241 | Yes | T | A |
| 13 | 87763670 | Yes | T | A |
| 13 | 88017138 | Yes | C | T |
| 13 | 88697850 | Yes | C | A |
| 13 | 88943911 | Yes | C | T |
| 13 | 89661680 | Yes | G | T |
| 13 | 90643059 | Yes | T | G |
| 13 | 91380475 | Yes | G | T |
| 13 | 95766579 | Yes | G | A |
| 14 | 106729548 | Yes | C | A |
| 14 | 23980848 | Yes | C | A |
| 14 | 25898992 | Yes | C | A |
| 14 | 28695331 | Yes | A | C |
| 14 | 29438815 | Yes | C | A |
| 14 | 47072639 | Yes | A | C |
| 14 | 47896926 | Yes | G | A |
| 14 | 48847319 | Yes | G | T |
| 14 | 51938150 | Yes | C | T |
| 14 | 76510398 | No | GT | G,TT |
| 14 | 82134585 | Yes | A | T |
| 14 | 83414219 | Yes | C | T |
| 14 | 87806426 | Yes | C | T |
| 14 | 96316510 | Yes | G | T |
| 15 | 102045836 | Yes | G | A |
| 15 | 22436191 | Yes | A | G |
| 15 | 24722504 | Yes | C | T |
| 15 | 25798544 | Yes | G | A |
| 15 | 25938642 | Yes | C | T |
| 15 | 70490762 | Yes | C | T |
| 15 | 75766255 | Yes | T | C |
| 15 | 77715276 | Yes | A | G |
| 15 | 95209654 | Yes | G | A |
| 16 | 1545435 | Yes | C | T |
| 16 | 18186958 | Yes | C | A |
| 16 | 25772589 | Yes | C | G |
| 16 | 27688502 | Yes | C | T |
| 16 | 28114143 | Yes | A | G |
| 16 | 31773014 | No | #N/A | #N/A |
| 16 | 3368718 | Yes | G | C |
| 16 | 34984555 | Yes | G | A |
| 16 | 48795508 | Yes | G | T |
| 16 | 49238327 | Yes | G | A |
| 16 | 52326430 | Yes | A | T |
| 16 | 5463986 | Yes | G | A |
| 16 | 60071187 | Yes | G | A |
| 16 | 62305237 | Yes | A | T |
| 16 | 63423040 | Yes | C | T |
| 16 | 65753775 | Yes | T | A |
| 16 | 6691633 | Yes | G | T |
| 16 | 70558147 | Yes | A | G |
| 16 | 83362861 | Yes | G | A |
| 17 | 10131181 | Yes | A | G |
| 17 | 13075762 | Yes | C | A |
| 17 | 21434416 | Yes | C | G |
| 17 | 39446715 | Yes | C | T |
| 17 | 49849301 | Yes | C | A |
| 17 | 5893063 | Yes | C | T |
| 17 | 64395578 | Yes | T | C |
| 17 | 70651401 | Yes | G | A |
| 17 | 73282938 | Yes | C | A |
| 18 | 14688237 | Yes | A | G |
| 18 | 29709626 | Yes | A | T |
| 18 | 35990989 | Yes | G | T |
| 18 | 39533132 | Yes | T | G |
| 18 | 46151443 | Yes | C | T |
| 18 | 48578438 | Yes | T | A |
| 18 | 54995268 | No | #N/A | #N/A |
| 18 | 56927520 | Yes | C | A |
| 18 | 67874726 | Yes | G | T |
| 18 | 69139353 | Yes | C | T |
| 18 | 71851399 | Yes | G | C |
| 18 | 77194654 | Yes | C | T |
| 19 | 11831410 | Yes | G | A |
| 19 | 13073880 | Yes | A | T |
| 19 | 1959983 | Yes | C | T |
| 19 | 24194443 | Yes | G | C |
| 19 | 29753852 | Yes | G | T |
| 19 | 30795213 | Yes | C | T |
| 19 | 31615967 | Yes | C | T |
| 19 | 37818487 | No | #N/A | #N/A |
| 19 | 46494625 | Yes | G | A |
| 19 | 51272098 | Yes | C | T |
| 19 | 54023925 | Yes | G | A |
| 19 | 54368378 | Yes | G | A |
| 19 | 54368379 | Yes | C | T |
| 19 | 55410446 | Yes | G | A |
| 2 | 100620585 | Yes | C | A |
| 2 | 102654398 | Yes | C | A |
| 2 | 107480628 | Yes | G | A |
| 2 | 107966476 | Yes | G | A |
| 2 | 108887438 | Yes | C | A |
| 2 | 1151479 | Yes | T | A |
| 2 | 117986148 | Yes | G | A |
| 2 | 123681866 | Yes | G | A |
| 2 | 132390960 | Yes | G | T |
| 2 | 132933876 | Yes | G | A |
| 2 | 137717364 | Yes | G | A |
| 2 | 137866429 | Yes | T | G |
| 2 | 141043701 | Yes | C | A |
| 2 | 149138046 | Yes | A | G |
| 2 | 151015695 | Yes | C | A |
| 2 | 15228066 | Yes | C | T |
| 2 | 156469721 | Yes | T | C |
| 2 | 159489974 | Yes | G | A |
| 2 | 170302449 | Yes | G | A |
| 2 | 17383144 | Yes | A | T |
| 2 | 175529970 | Yes | C | T |
| 2 | 177419158 | Yes | G | A |
| 2 | 1793969 | Yes | G | A |
| 2 | 185713930 | Yes | T | A |
| 2 | 194930313 | Yes | G | C |
| 2 | 195667273 | Yes | G | A |
| 2 | 200455800 | Yes | C | T |
| 2 | 210389779 | Yes | G | A |
| 2 | 214333099 | Yes | C | A |
| 2 | 215167616 | Yes | T | C |
| 2 | 218752092 | Yes | C | T |
| 2 | 221403747 | Yes | G | A |
| 2 | 221867294 | Yes | A | G |
| 2 | 230987206 | Yes | C | T |
| 2 | 240196595 | Yes | G | T |
| 2 | 35901623 | Yes | G | A |
| 2 | 35981902 | Yes | C | T |
| 2 | 36443679 | Yes | C | T |
| 2 | 42463791 | Yes | C | T |
| 2 | 48227618 | Yes | T | G |
| 2 | 50996088 | Yes | T | A |
| 2 | 51036672 | Yes | T | G |
| 2 | 52410090 | Yes | G | A |
| 2 | 69820777 | Yes | A | G |
| 2 | 78139135 | Yes | A | C |
| 2 | 78232835 | Yes | C | A |
| 2 | 84009712 | Yes | A | G |
| 2 | 99845188 | Yes | C | A |
| 20 | 11349749 | Yes | A | G |
| 20 | 12578989 | Yes | G | A |
| 20 | 34988121 | Yes | C | T |
| 20 | 42697100 | Yes | G | A |
| 20 | 52022310 | Yes | G | A |
| 20 | 53497524 | Yes | G | T |
| 20 | 53831019 | Yes | C | G |
| 20 | 59122074 | Yes | A | G |
| 20 | 59762970 | Yes | A | T |
| 21 | 23845788 | Yes | G | A |
| 21 | 24379137 | Yes | C | A |
| 21 | 25951524 | Yes | G | A |
| 21 | 39423946 | Yes | T | C |
| 21 | 43746189 | Yes | G | A |
| 21 | 45974068 | Yes | T | A |
| 22 | 28034479 | Yes | G | A |
| 22 | 32957641 | Yes | C | T |
| 22 | 38256655 | Yes | C | A |
| 22 | 44367335 | Yes | G | A |
| 22 | 47586479 | Yes | G | A |
| 22 | 48126819 | Yes | C | T |
| 22 | 51109824 | Yes | C | T |
| 3 | 101793395 | Yes | G | A |
| 3 | 10237999 | Yes | A | T |
| 3 | 104125697 | Yes | G | C |
| 3 | 10835796 | Yes | C | G |
| 3 | 109960669 | Yes | T | C |
| 3 | 1102211 | Yes | C | T |
| 3 | 110614435 | Yes | G | A |
| 3 | 116625346 | Yes | C | A |
| 3 | 124273108 | Yes | C | G |
| 3 | 124483249 | Yes | G | C |
| 3 | 130262179 | Yes | C | A |
| 3 | 131747033 | Yes | C | T |
| 3 | 134555943 | Yes | C | T |
| 3 | 141327236 | Yes | C | A |
| 3 | 145251251 | Yes | C | T |
| 3 | 147576669 | Yes | G | T |
| 3 | 15525552 | Yes | T | C |
| 3 | 163642970 | Yes | G | A |
| 3 | 164222721 | Yes | G | A |
| 3 | 164713933 | Yes | G | A |
| 3 | 166079986 | Yes | T | A |
| 3 | 170371610 | Yes | C | T |
| 3 | 18168560 | Yes | G | A |
| 3 | 182529805 | Yes | C | T |
| 3 | 186697776 | Yes | C | T |
| 3 | 19133133 | Yes | G | A |
| 3 | 191452623 | Yes | C | T |
| 3 | 195653781 | No | #N/A | #N/A |
| 3 | 21850915 | Yes | G | T |
| 3 | 24247783 | Yes | #N/A | #N/A |
| 3 | 29841231 | Yes | T | C |
| 3 | 53870279 | Yes | C | T |
| 3 | 5560596 | Yes | T | A |
| 3 | 6359197 | Yes | G | C |
| 3 | 65150590 | Yes | C | T |
| 3 | 65437969 | Yes | T | C |
| 3 | 69417779 | Yes | A | T |
| 3 | 73578475 | Yes | G | A |
| 3 | 7760915 | Yes | C | A |
| 3 | 77752655 | Yes | G | A |
| 3 | 78301834 | Yes | A | G |
| 3 | 84590333 | Yes | G | A |
| 3 | 87615315 | Yes | T | A |
| 3 | 90093839 | Yes | A | G |
| 4 | 105210715 | Yes | C | T |
| 4 | 105771830 | Yes | C | T |
| 4 | 116988184 | Yes | G | A |
| 4 | 121062576 | Yes | G | A |
| 4 | 121267820 | Yes | T | G |
| 4 | 122845863 | Yes | T | G |
| 4 | 131429584 | Yes | C | T |
| 4 | 131521084 | Yes | T | A |
| 4 | 131565643 | Yes | A | G |
| 4 | 136883134 | Yes | C | A |
| 4 | 137223054 | Yes | G | T |
| 4 | 141430215 | Yes | C | T |
| 4 | 143630538 | Yes | G | A |
| 4 | 144564154 | Yes | C | T |
| 4 | 149451685 | Yes | A | T |
| 4 | 154921582 | Yes | A | G |
| 4 | 160238557 | Yes | G | A |
| 4 | 16405139 | Yes | T | G |
| 4 | 164791887 | Yes | C | T |
| 4 | 164939381 | Yes | T | G |
| 4 | 167559756 | Yes | T | G |
| 4 | 168406993 | Yes | G | T |
| 4 | 169051729 | Yes | A | G |
| 4 | 170131829 | Yes | G | A |
| 4 | 170900891 | Yes | G | C |
| 4 | 174354602 | Yes | C | T |
| 4 | 177810095 | Yes | T | A |
| 4 | 182464840 | Yes | C | A |
| 4 | 189974174 | Yes | G | T |
| 4 | 19040755 | Yes | T | A |
| 4 | 26522531 | Yes | T | C |
| 4 | 27615114 | Yes | C | T |
| 4 | 29271645 | Yes | G | T |
| 4 | 29332629 | Yes | T | C |
| 4 | 32674148 | Yes | C | T |
| 4 | 33349776 | Yes | T | A |
| 4 | 33372395 | Yes | A | T |
| 4 | 43851763 | Yes | G | A |
| 4 | 45844938 | Yes | A | T |
| 4 | 47094927 | Yes | G | A |
| 4 | 53118559 | Yes | A | G |
| 4 | 58337699 | Yes | T | C |
| 4 | 5919971 | Yes | G | C |
| 4 | 60376453 | Yes | T | C |
| 4 | 61381784 | Yes | T | G |
| 4 | 63743899 | Yes | G | C |
| 4 | 64070881 | Yes | T | A |
| 4 | 65051960 | Yes | C | A |
| 4 | 65051962 | Yes | C | A |
| 4 | 65148226 | Yes | T | G |
| 4 | 66916687 | Yes | G | T |
| 4 | 80451158 | Yes | G | T |
| 4 | 82550138 | Yes | T | A |
| 4 | 85162597 | Yes | C | T |
| 4 | 92690249 | Yes | G | A |
| 4 | 97716391 | Yes | C | T |
| 4 | 98034563 | Yes | T | A |
| 4 | 98743853 | Yes | G | A |
| 4 | 99480116 | Yes | T | A |
| 5 | 104518420 | Yes | C | T |
| 5 | 112539816 | Yes | G | C |
| 5 | 11792010 | Yes | A | T |
| 5 | 119676379 | Yes | T | A |
| 5 | 122828075 | Yes | G | A |
| 5 | 125305112 | Yes | T | C |
| 5 | 127310968 | Yes | C | T |
| 5 | 136162747 | Yes | G | A |
| 5 | 145996883 | Yes | T | C |
| 5 | 146729397 | Yes | G | A |
| 5 | 153019851 | Yes | C | T |
| 5 | 153938181 | Yes | C | T |
| 5 | 15400433 | Yes | C | A |
| 5 | 154798740 | Yes | G | T |
| 5 | 159177218 | Yes | G | T |
| 5 | 160610822 | Yes | T | G |
| 5 | 162976623 | Yes | T | C |
| 5 | 167264050 | Yes | A | G |
| 5 | 169948406 | Yes | A | C |
| 5 | 29416842 | Yes | C | G |
| 5 | 30961029 | Yes | G | A |
| 5 | 44047485 | Yes | C | A |
| 5 | 44047490 | Yes | G | T |
| 5 | 44369620 | Yes | C | T |
| 5 | 4630333 | Yes | T | C |
| 5 | 50394596 | Yes | T | G |
| 5 | 50975661 | Yes | G | A |
| 5 | 51028135 | Yes | C | T |
| 5 | 58230047 | Yes | C | G |
| 5 | 60652809 | Yes | G | A |
| 5 | 61568543 | Yes | G | T |
| 5 | 75070604 | Yes | T | A |
| 5 | 7532565 | Yes | C | G |
| 5 | 76124493 | Yes | A | T |
| 5 | 84466498 | Yes | G | T |
| 5 | 85895640 | Yes | T | A |
| 5 | 87884834 | Yes | C | A |
| 5 | 89015268 | Yes | A | G |
| 5 | 8958583 | Yes | T | A |
| 5 | 89666900 | Yes | G | A |
| 5 | 99763455 | Yes | T | C |
| 6 | 104372462 | Yes | A | G |
| 6 | 116696900 | Yes | C | T |
| 6 | 119720336 | Yes | T | G |
| 6 | 123611009 | Yes | G | T |
| 6 | 127524508 | Yes | G | T |
| 6 | 141669541 | Yes | A | G |
| 6 | 141695502 | Yes | T | G |
| 6 | 144159636 | Yes | C | T |
| 6 | 148044200 | Yes | C | T |
| 6 | 167832727 | Yes | A | C |
| 6 | 18981715 | Yes | G | C |
| 6 | 21032904 | Yes | C | G |
| 6 | 21077120 | Yes | C | G |
| 6 | 22051742 | Yes | G | T |
| 6 | 3317706 | Yes | G | C |
| 6 | 55353772 | Yes | G | A |
| 6 | 62195820 | Yes | G | T |
| 6 | 65974058 | Yes | T | C |
| 6 | 66175981 | Yes | C | T |
| 6 | 67703856 | Yes | A | T |
| 6 | 82143420 | Yes | G | A |
| 6 | 91020872 | Yes | G | A |
| 6 | 94337688 | Yes | A | G |
| 6 | 9898321 | Yes | T | C |
| 6 | 9898333 | Yes | G | C |
| 7 | 106060713 | Yes | G | T |
| 7 | 108105873 | Yes | C | T |
| 7 | 136742792 | Yes | G | C |
| 7 | 144171676 | Yes | A | G |
| 7 | 145122310 | Yes | C | T |
| 7 | 145957657 | Yes | A | T |
| 7 | 146492548 | Yes | G | T |
| 7 | 147744928 | Yes | G | A |
| 7 | 147878170 | Yes | C | G |
| 7 | 153082728 | Yes | T | C |
| 7 | 155824366 | Yes | A | G |
| 7 | 17625224 | Yes | G | C |
| 7 | 19371977 | Yes | T | G |
| 7 | 2256785 | Yes | C | G |
| 7 | 39191208 | Yes | T | A |
| 7 | 40835867 | Yes | C | A |
| 7 | 41021059 | Yes | T | C |
| 7 | 43173850 | Yes | G | A |
| 7 | 49617010 | Yes | G | A |
| 7 | 54100652 | Yes | C | T |
| 7 | 6272829 | Yes | C | A |
| 7 | 70708908 | Yes | A | C |
| 7 | 73876676 | Yes | A | C |
| 7 | 79154392 | Yes | G | A |
| 7 | 82380633 | Yes | G | A |
| 7 | 83550706 | Yes | C | G |
| 7 | 89881547 | Yes | C | A |
| 7 | 9497310 | Yes | C | A |
| 8 | 100329974 | Yes | A | T |
| 8 | 104367943 | Yes | T | A |
| 8 | 105016894 | Yes | C | T |
| 8 | 106838897 | Yes | T | C |
| 8 | 110689781 | Yes | G | A |
| 8 | 110981676 | Yes | T | C |
| 8 | 11176892 | Yes | G | T |
| 8 | 114788070 | Yes | G | A |
| 8 | 115861860 | Yes | T | C |
| 8 | 117286756 | Yes | A | G |
| 8 | 126954162 | Yes | T | A |
| 8 | 131333555 | Yes | A | T |
| 8 | 137400891 | Yes | C | A |
| 8 | 138124461 | Yes | G | A |
| 8 | 142410720 | Yes | G | A |
| 8 | 144140505 | Yes | G | A |
| 8 | 15842720 | Yes | G | C |
| 8 | 19202942 | Yes | G | A |
| 8 | 2361814 | Yes | G | A |
| 8 | 26516789 | Yes | A | G |
| 8 | 34049464 | Yes | C | G |
| 8 | 55830938 | Yes | A | C |
| 8 | 57667175 | Yes | T | C |
| 8 | 58919918 | Yes | T | C |
| 8 | 59419315 | Yes | G | T |
| 8 | 61450653 | Yes | C | G |
| 8 | 62972325 | Yes | A | G |
| 8 | 64724856 | Yes | G | T |
| 8 | 66090168 | Yes | C | T |
| 8 | 72178967 | Yes | G | T |
| 8 | 76653780 | Yes | A | G |
| 8 | 78632217 | Yes | C | T |
| 8 | 78788167 | Yes | C | T |
| 8 | 81939357 | Yes | C | T |
| 8 | 83132659 | Yes | C | G |
| 8 | 87930503 | Yes | T | C |
| 8 | 91571842 | Yes | C | A |
| 8 | 93527163 | Yes | G | T |
| 9 | 106820320 | Yes | C | T |
| 9 | 110548557 | Yes | A | C |
| 9 | 110948342 | Yes | C | T |
| 9 | 118876506 | Yes | A | T |
| 9 | 131122810 | Yes | G | A |
| 9 | 18234501 | Yes | A | G |
| 9 | 21969488 | Yes | A | C |
| 9 | 28394129 | Yes | C | T |
| 9 | 28791686 | Yes | C | G |
| 9 | 29034163 | Yes | G | A |
| 9 | 29141215 | Yes | C | A |
| 9 | 30148095 | Yes | T | A |
| 9 | 32406942 | Yes | C | A |
| 9 | 43647126 | Yes | T | A |
| 9 | 71868478 | Yes | C | T |
| 9 | 76590210 | No | #N/A | #N/A |
| 9 | 76590227 | No | #N/A | #N/A |
| 9 | 80291817 | Yes | T | C |
| 9 | 82464856 | Yes | C | A |
| 9 | 8403405 | Yes | G | T |
| 9 | 8543648 | Yes | C | T |
| 9 | 87247996 | Yes | C | T |

**Table S3.** All identified base pair substitutions that affect protein coding regions of the genome

| Individual | Tissue | Chromosome | Position | Mutation type | Consequence | Gene strand | Gene name |
| --- | --- | --- | --- | --- | --- | --- | --- |
| w22_1 | Liver | chr19 | 17435766 | T>A | nonsense | - | ANO8 |
| w22_2 | Liver | chr2 | 39406378 | G>T | nonsynonymous | - | CDKL4 |
| w15 | Intestine | chr11 | 65404068 | G>A | nonsynonymous | + | PCNX3 |
| w15 | Liver | chr16 | 1545435 | C>T | nonsynonymous | + | TELO2 |
| w17 | Intestine | chr1 | 27210699 | A>G | nonsynonymous | - | GPN2 |
| w17 | Liver | chr3 | 124483249 | G>C | nonsynonymous | - | ITGB5 |

